# The anterior paired lateral neuron normalizes odour-evoked activity at the mushroom body calyx

**DOI:** 10.1101/2021.09.20.461071

**Authors:** Luigi Prisco, Stephan Hubertus Deimel, Hanna Yeliseyeva, André Fiala, Gaia Tavosanis

## Abstract

To identify and memorize discrete but similar environmental inputs, the brain needs to distinguish between subtle differences of activity patterns in defined neuronal populations. The Kenyon cells of the *Drosophila* adult mushroom body (MB) respond sparsely to complex olfactory input, a property that is thought to support stimuli discrimination in the MB. To understand how this property emerges, we investigated the role of the inhibitory anterior paired lateral neuron (APL) in the input circuit of the MB, the calyx. Within the calyx, presynaptic boutons of projection neurons (PNs) form large synaptic microglomeruli (MGs) with dendrites of postsynaptic Kenyon cells (KCs). Combining EM data analysis and *in vivo* calcium imaging, we show that APL, via inhibitory and reciprocal synapses targeting both PN boutons and KC dendrites, normalizes odour-evoked representations in MGs of the calyx. APL response scales with the PN input strength and is regionalized around PN input distribution. Our data indicate that the formation of a sparse code by the Kenyon cells requires APL-driven normalization of their MG postsynaptic responses. This work provides experimental insights on how inhibition shapes sensory information representation in a higher brain centre, thereby supporting stimuli discrimination and allowing for efficient associative memory formation.

## 1 Introduction

Every day we are challenged to navigate through a complex and variable environment, often characterized by similar stimuli combined in different ways. Yet, our brain excels in assessing if, and how, the current experience is different or similar to a previously encountered one. The ability to discriminate across stimuli is achieved by minimizing the overlap between patterns of neuronal activity through a process defined as “pattern separation” (Santoro 2013). This conserved property is intrinsic to diverse circuits such as the mammalian cerebellum, the dentate gyrus and the *Drosophila* mushroom body (MB) (Cayco-Gajic and Silver 2019). In the current models, all the aforementioned circuits support pattern separation by utilizing different degree of inhibitory mechanisms. (Tyrrell and Willshaw 1992; Schweighofer, Doya, and Lay 2001; Sahay, Wilson, and Hen 2011; Cayco-Gajic, Clopath, and Silver 2017; Litwin-Kumar et al. 2017). Experimental evidence in support of these inhibitory circuits has been described over the years (Vos, Volny-Luraghi, and de Schutter 1999; Duguid et al. 2015; Inada, Tsuchimoto, and Kazama 2017; Parnas et al. 2013; Olsen, Bhandawat, and Wilson 2010; A. C. Lin et al. 2014), however, the mechanism by which inhibition contributes to pattern separation is not yet fully understood, often due to technical limitations.

With an extended genetic toolkit and a brain of only ∼100,000 neurons (Raji and Potter 2021; Alivisatos et al. 2012) largely reconstructed at the EM Level (Zheng et al. 2018; F. Li et al. 2020a), *Drosophila* represents an attractive system to provide experimental evidence on the mechanisms behind pattern separation. The fly MB receives mainly olfactory input, though optical, temperature and humidity information is also represented (Marin et al. 2020; Frank et al. 2015; J. Li et al. 2020). The MB is required for memory formation and retrieval (Heisenberg et al. 1985; de Belle and Heisenberg 1994; Dubnau et al. 2001; S. E. McGuire, Le, and Davis 2001; Aso et al. 2014). Within the MB input region, in the main calyx, olfactory projection neurons (PNs) deliver sensory information from 51 distinct olfactory glomeruli (Grabe et al. 2016; Bates et al. 2020) to ∼2,000 Kenyon cells (KCs) of the MB (Aso et al. 2009), for an expansion ratio of 40 (Litwin-Kumar et al. 2017). In the calyx, PNs synapse onto KCs via complex synaptic structures known as microglomeruli (MGs) (Yasuyama, Meinertzhagen, and Schürmann 2002a; Leiss et al. 2009a). At each MG, a single central PN bouton is enwrapped by, on average, 13 claw-like dendritic terminals of as many different KCs (Davi D. Bock, personal communication). KCs integrate inputs in a combinatorial manner, with each KC receiving input from 6-8 PNs, on average (Butcher et al. 2012; Zheng et al. 2020; F. Li et al. 2020b; Turner, Bazhenov, and Laurent 2008), of which more than half need to be coactive to elicit spikes (Gruntman and Turner 2013; Inada, Tsuchimoto, and Kazama 2017). As a result, while PN odour-evoked activity is broadly tuned (Perez-Orive et al. 2002; Bhandawat et al. 2007), odour representation is sparse and decorrelated at the KCs layer (Honegger, Campbell, and Turner 2011; Turner, Bazhenov, and Laurent 2008; Campbell et al. 2013b; Perez-Orive et al. 2002), therefore reducing overlap between stimuli representation and allowing for better discriminability (Kanerva 1988; Cayco-Gajic, Clopath, and Silver 2017; Olshausen and Field 2004). In addition to sparse PN:KC connectivity and KCs high input threshold, inhibition is required to reduce the overlap among odour representations in the *Drosophila* MB (A. C. Lin et al. 2014; Lei et al. 2013). At the MB, inhibition is provided by the GABAergic anterior paired lateral (APL) neuron, which innervates both the calyx and the lobes of the MB (Liu and Davis 2009; Pitman et al. 2011; Aso et al. 2014). APL responds to odours with depolarization and calcium influx (Liu and Davis 2009; Papadopoulou et al. 2011). Importantly, blocking APL output disrupts the KCs sparse odour representation and impairs learned discrimination of similar odours, pointing to its critical role in the process (A. C. Lin et al. 2014; Lei et al. 2013). APL is suggested to regulate sparse coding by participating in a closed feedback loop with the MB, similarly to its homolog giant GABAergic neuron (GGN) in the locust (Papadopoulou et al. 2011; A. C. Lin et al. 2014; Litwin-Kumar et al. 2017). However, APL is both pre- and post- synaptic to PNs and KCs in the adult calyx (Yasuyama, Meinertzhagen, and Schürmann 2002b; Wu et al. 2013; Baltruschat et al. 2021). Additionally, APL response to localized stimuli is spatially restricted (Amin et al. 2020). In particular, APL branches at the MB lobes and the ones in the calyx appear to represent two separate compartments (Amin et al. 2020), suggesting a possible distinct role of APL inhibition in these two different compartments. Hence, the mechanisms by which APL modulates sparse coding and its involvement in the process of pattern separation are still unclear. In the present work, we challenge the concept of a broad feedback inhibition to the MB calyx by APL with primary experimental data. In particular, we focused on the APL processes within the MB calyx and set out to identify the role of GABAergic inhibition at the PN:KC synaptic layer. Taking advantage of recently released EM datasets (Scheffer et al. 2020; Zheng et al. 2018), we report the complex synaptic interaction of APL with PNs and KCs within the MGs of the MB calyx. Next, via *in vivo* calcium imaging in the calyx, we explored the role of APL inhibition onto MGs by recording the odour-evoked activity of APL, PN boutons and KC dendritic claws. Our results indicate that APL acts as a normalizer of postsynaptic responses to olfactory inputs in the MGs of the mushroom body calyx, an idea that we confirmed by blocking the output of APL. Additionally, via volumetric calcium imaging, we addressed the locality of APL activation in the calyx and found that it is odour-specific. We suggest that the normalization of postsynaptic MG responses by APL is essential to determine the key property of KCs to respond only to the coincident input of PNs to multiple claws, allowing for an elevated stimulus discriminability.

## 2 Results

### 2.1 APL is an integral part of the microcircuit within microglomeruli in the MB calyx

To better understand the role of GABAergic inhibition at the MB calyx, we investigated APL involvement into the calycal microcircuits with the highest resolution available. The APL of adult *Drosophila* innervates extensively all compartments of the mushroom body, including calyx, lobes and pedunculus (Liu and Davis 2009). Moreover, the neuron appears to be non-polarized in the adult, with strong expression of both pre- and post-synaptic markers in all compartments (Wu et al. 2013). However, little is known regarding the detailed connectivity between APL and the cell types constituting the mushroom body.

Taking advantage of emerging electron microscopy (EM) datasets covering a full adult fly brain (FAFB, (Zheng et al. 2018)) or a large fraction of it (Hemibrain, (Scheffer et al. 2020)), we examined the distribution of synaptic contacts between APL, PNs and KCs, the major cell types constituting the MGs of the mushroom body calyx (Leiss et al. 2009b; Yasuyama, Meinertzhagen, and Schürmann 2002a; Baltruschat et al. 2021) (Fig 1A). We recently reconstructed an entire MG in the FAFB dataset, starting from a PN-bouton of the DA1 glomerulus and tracing all its pre- and post-synaptic partners (Baltruschat et al. 2021). Here, we focused on the synaptic connections involving APL. We found APL to be highly involved in the MG structure, with pre- and post- synaptic contacts with both KC dendrites and PN boutons (Fig S1A) (Baltruschat et al. 2021). Many of those synapses were polyadic, displaying typical configurations within that specific MG (described in Fig S1B). To verify whether such features were specific to the DA1 MG reconstructed in Baltruschat et al. (2021) or common, we exploited the Hemibrain EM dataset (Scheffer et al. 2020) and extracted all calycal connections from and to APL with either KCs or PNs. Out of the 136 PNs reported innervating the main calyx (F. Li et al. 2020b), 126 made and received synapses with APL (full list of PNs and APL interactions available at: Mendeley data link will be available upon publication). To reveal the localization of these synapses, we rendered 3D graphs of single PNs derived from the Hemibrain dataset (Scheffer et al. 2020) and mapped the synapses that they receive from APL within the MB calyx (see Material and Methods for details). Most of the APL-to-PN connections were localized on PN boutons (84 ± 2%, mean ± SEM, of the total synapses received by each PN localised on boutons), demonstrating that the majority of APL-PN interactions happens at MGs (Fig 1B, all images available at: Mendeley data link will be available upon publication). Additionally, we found a positive correlation between the number of synapses made by the APL towards a specific PN and the reciprocal synapses formed by that PN onto APL (Fig 1D).

**Fig 1.**
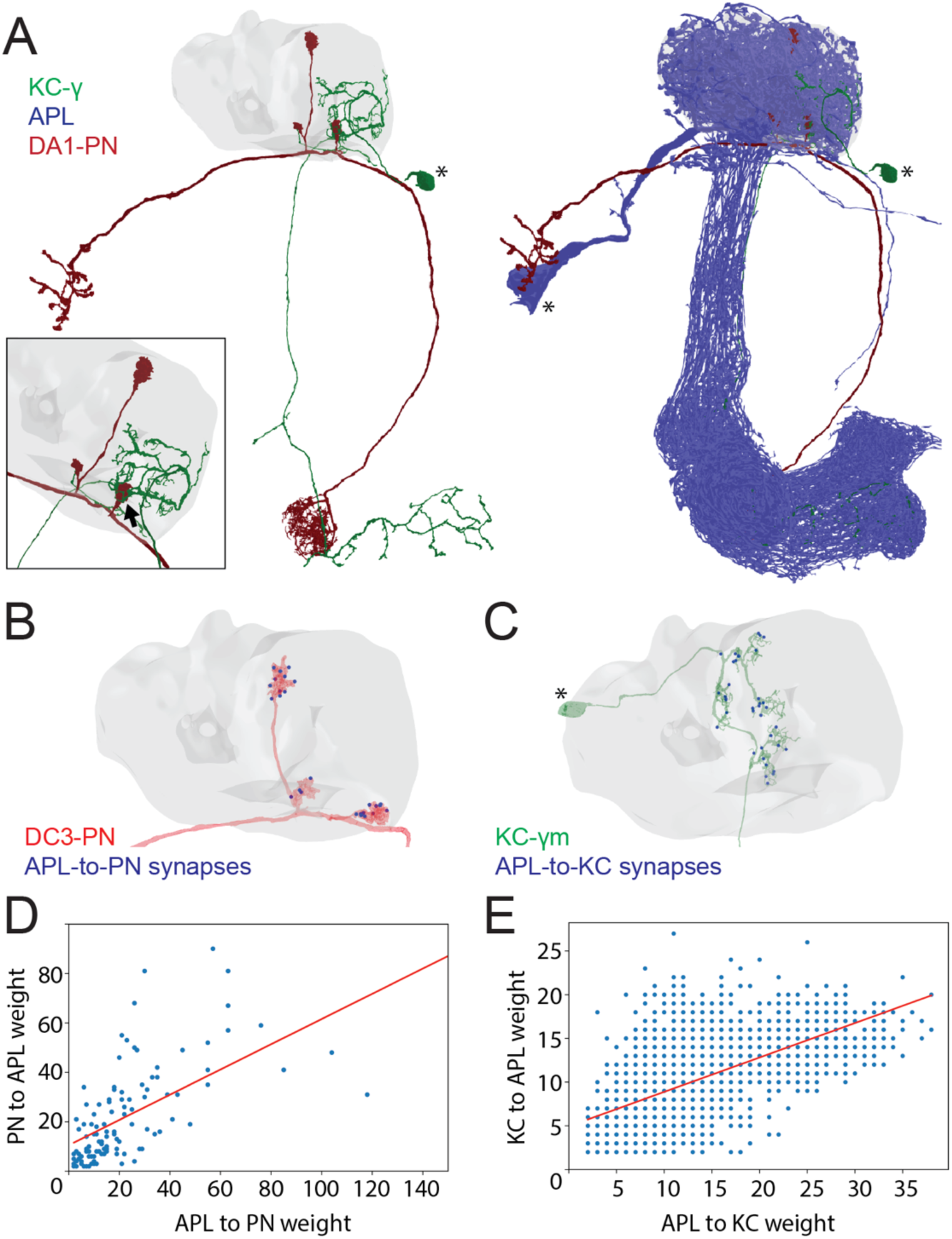
APL participates in the MG microcircuit with reciprocal synapses. **(A)** Left: example of a PN (red) sending collateral boutons into the MB calyx (grey volume), where it connects onto KCs claws via synaptic microglomeruli. For simplicity, only 1 KC is visualised here (green). Bottom-left box: magnification of a PN bouton interacting with a KC claw (black arrow). Right: APL (blue) innervates the entire MB including lobes, peduncle and the calyx. Asterisks indicate cell bodies. **(B)** Visualisation of APL synapses (blue dots) onto a PN 3D mesh within the MB calyx. Most connections are localised on PN boutons. **(C)** Localization of APL synapses (blue) on a KC 3D mesh within the MB calyx. While most are localised on dendritic claws, some connections along dendritic branches could be seen as well (see also S1C). The cell body is marked by an asterisk. **(D)** Correlation between the number of PN-to-APL reciprocal synapses (r^2^=0.63) and KC-to-APL ones **(E)** (r^2^=0.60). The correlation was calculated among the entire synaptic weight that individual PNs or KCs had with APL. All 3D plots were created via the Neuprint-python package (see Materials & Methods).

Of notice, most of the PNs not connecting to the APL within the main calyx were already described as non-olfactory PNs (Marin et al. 2020), and they all seemed to extend most of their terminals elsewhere, with little to no branches in the main calyx (Fig S1D). Similarly, of the 1919 KCs present in the dataset, 1871 displayed interactions with APL. Mapping APL synapses onto single KC meshes (Fig 1C, all images available at: Mendeley data link will be available upon publication) showed a majority of connections on KC claws. However, we noticed inhibitory synapses along KC dendrites as well. The KCs constituting the MB are divided in 3 major classes based on their axonal projections: *γ*, *α*/*β*, *α*’/*β*’ (Crittenden et al. 1998; Lee, Lee, and Luo 1999). We found a difference in the spatial distribution of APL synapses depending on the KC type, suggesting that APL inhibition might have a different impact on different KC types. In particular, APL synapses onto *α*/*β* KCs were significantly less localised on claw-like dendritic terminals and more distributed along KC dendritic branches (Fig S1C, n=210 (70 per KC type, randomly selected), p<0.0001, Unpaired ANOVA with multiple comparisons). The KCs not interacting with APL displayed a rather atypical structure, with extensive dendritic arborization just outside of the main calyx rather than within (Fig S1E). As in the case of PNs, the number of KC-to-APL synapses positively correlated with the APL-to-KC synapse number (Fig 1E). In conclusion, EM dataset analysis revealed a large involvement of APL in the calycal circuitry, with reciprocal connections to the vast majority of PNs and KCs. APL involvement in the MG structure as reported in Baltruschat et al (2021) might be thus generalized to potentially all MGs of the main calyx.

### 2.2 In the calyx, APL displays different response levels to different odours

The analysis of the EM data provided structural evidence for possible feedforward and feedback circuits between APL, PNs and KCs in the MB calyx (Fig S1A-B). To explore the functional role of APL in calycal MGs, we performed *in vivo* functional imaging experiments by expressing the calcium indicator GCaMP6m (Chen et al. 2013b) specifically in APL via the APL intersectional *NP2631-GAL4, GH146-FLP* (APLi) driver (A. C. Lin et al. 2014; Mayseless et al. 2018) and recorded odour-evoked activity in the calyx (Fig 2A, see Materials and Methods). Flies were stimulated with 5 second puffs of odours diluted 1:100 in mineral oil and exposed to sequences of 2 odours starting with 4- methylcyclohexanol (Mch) and 3-octanol (Oct), presented in a randomized fashion. Odour-elicited calcium transients in APL were detectable in the calyx (Fig 2B,E). Interestingly, we observed a clear difference in the GCaMP fluorescence levels, measured as *Δ*F/F_0_ over the entire calycal region innervated by APL (see also Material and Methods) in response to the two odour stimulations, with Oct eliciting a stronger APL response (Fig 2C-D, n=10, p=0.002, Wilcoxon matched-pairs test). To extend this observation, we exposed flies also to ẟ-Decalactone (ẟ-DL), an odour reported to elicit the least overall activity in ORNs (Hallem and Carlson 2006a). Similarly, we measured a difference between the strength of the response to Oct compared to ẟ-DL (Fig 2F-G, n=10, p<0.0001, paired t test). Moreover, the gap between the ẟ-DL signal peak and the Oct one was higher compared to the Mch vs Oct group (Δ(Oct-Mch) = 45 ± 27%; Δ(Oct-ẟ-DL) = 76 ± 30%, n=10, p=0.0234, unpaired t-test with Welch’s correction), suggesting that APL is able to provide a variable, odour-tuned inhibition to MGs of the MB calyx.

**Figure 2.**
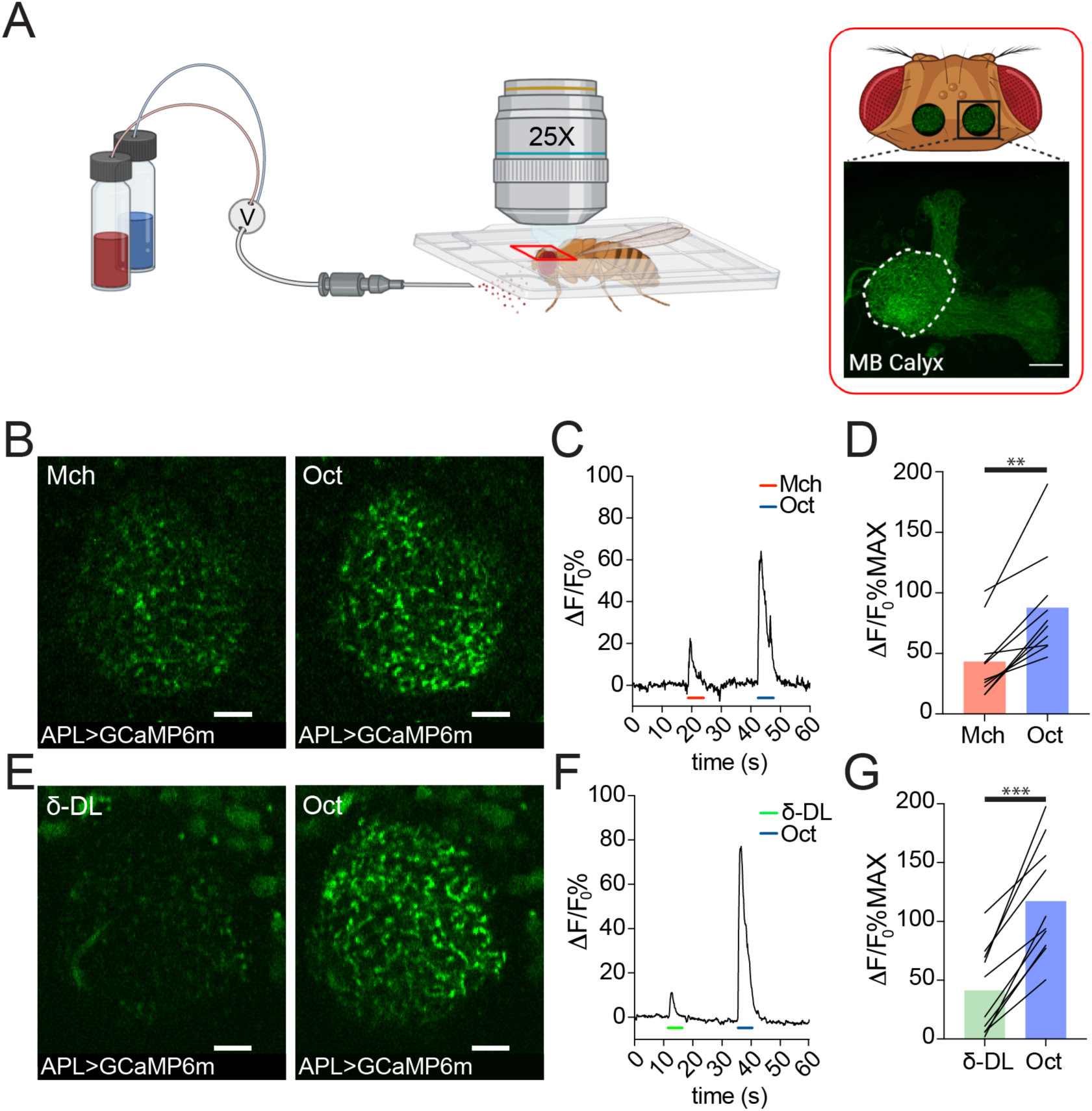
APL responds to odours with variable calcium transients. **(A)** Schematic view of the two-photon *in vivo* imaging setup. Scale bar = 20μm. **(B)** Example of APL response to Mch or Oct in the calyx of *APLi- GAL4>UAS-GCaMP6m* flies. Scale bar = 10μm. **(C)** Fluorescence intensity over time for the fly showed in (B). **(D)** APL showed higher intracellular calcium transients in response to Oct compared to Mch. n=10, p=0.002, Wilcoxon matched-pairs test. **(E)** Example of APL GCaMP6m response to *δ*-DL or Oct. Scale bar = 10μm. **(F)** Fluorescence intensity over time for the fly showed in (E). **(G)** APL peak response comparison for the *δ*-DL vs Oct odours sequence. n=10, p<0.0001, paired t test. Odours were diluted 1:100, bars indicate means.

### 2.3 The response to different odours is highly variable in PNs, but more homogeneous in KC dendrites

To investigate the origin and the consequences of the observed difference in APL response at the MB calyx, we performed functional imaging experiments targeting the other two cell types participating in the microglomerular structure: PNs and KCs (Leiss et al. 2009b; Yasuyama, Meinertzhagen, and Schürmann 2002a). Odors are detected by a large set of olfactory receptor neurons (ORNs) expressing chemically-tuned odorant receptors (Clyne et al. 1999; Hallem and Carlson 2006a). ORNs project to the 51 distinct olfactory glomeruli in the adult antennal lobe (AL) in a stereotyped manner, with ORNs expressing the same odorant receptor projecting to the same glomerulus (Q. Gao, Yuan, and Chess 2000; Vosshall, Wong, and Axel 2000; Grabe et al. 2016). Within glomeruli, ORNs synapse onto second-order neurons, the PNs, which deliver odour information to higher brain regions such as the MB and the lateral horn (R. F. Stocker et al. 1990). To investigate whether odour-evoked activity in PNs reflected the differences in strength observed in APL, we expressed GCaMP6m in PNs via the generic PN-Gal4 driver *GH146* (Berdnik et al. 2008), and imaged PN dendrites in the antennal lobe. Flies were exposed to Mch or Oct as described for the APL imaging experiments, and the average peak among the responding glomeruli per brain was used as a general indicator of the total input transmitted by PNs. Imaging was performed on a single optical section of the antennal lobe, and only the glomeruli that could be unequivocally identified among all tested animals were taken into consideration for the analysis. While the number of responding glomeruli was similar between the two tested odours (Fig S2, also shown on a larger number of AL glomeruli in (Barth et al. 2014)), the overall calcium transient was higher when flies were exposed to Oct (Fig S3A-B, n=10, p=0.002, Wilcoxon matched-pairs test), suggesting that the main source of difference was represented by the degree of PNs activation rather than an additional/decreased number of active neurons. Likewise, a strong difference could be measured when flies were exposed to the ẟ-DL/Oct odours sequence (Fig S3C-D, n=10, p<0.0001, paired t-test), resembling the differences in APL activation detected at the MB calyx.

To address whether this odour-dependent variability in PN dendrites activity is still detectable within the collateral boutons in the MB calyx, we expressed the presynaptically localized GCaMP3 transgene *UAS-Syp::GCaMP3* (Pech et al. 2015) in PNs and recorded odour-evoked activity in PN boutons of the MB calyx. We exposed flies to Mch or Oct, and calculated per calyx the average peak response among the boutons showing calcium transients. While the number of active boutons did not change between Mch and Oct stimulations (Fig 3D, n=10, p=0.689, paired t-test), the average boutons response was higher when flies were exposed to Oct (Fig 3B, n=10, p=0.0002, paired t-test). Furthermore, plotting the frequency distribution of all boutons activity peaks measured during these experiments showed a clear shift towards higher values of the entire Oct-responding population (Fig 3C, n=10, p<0.0001, Kolmogorov-Smirnov test). Hence, the difference shown in Fig 3B was not just due to a very high response of a few boutons, but rather to an overall increase in PN boutons activation levels across stimuli . Taken together, these experimental data suggested that the APL neuron activation scales with PN inputs strength. To further extend the correlation between PNs and APL activity in a systematic way, we measured the APL calcium transient levels in response to odours contained in a ORNs response database (Hallem and Carlson 2006b) and plotted it against their PN spikes value obtained via the experimentally supported equation described in Olsen et al. (Olsen, Bhandawat, and Wilson 2010; Parnas et al. 2013). We observed a positive linearity between the two variables (Fig S3E, Pearson r=0.99), hence supporting our experimental data described above. In conclusion, these experiments demonstrated that the overall level of PN activation varies with different odours and correlates with the APL response within the MB calyx.

**Fig 3.**
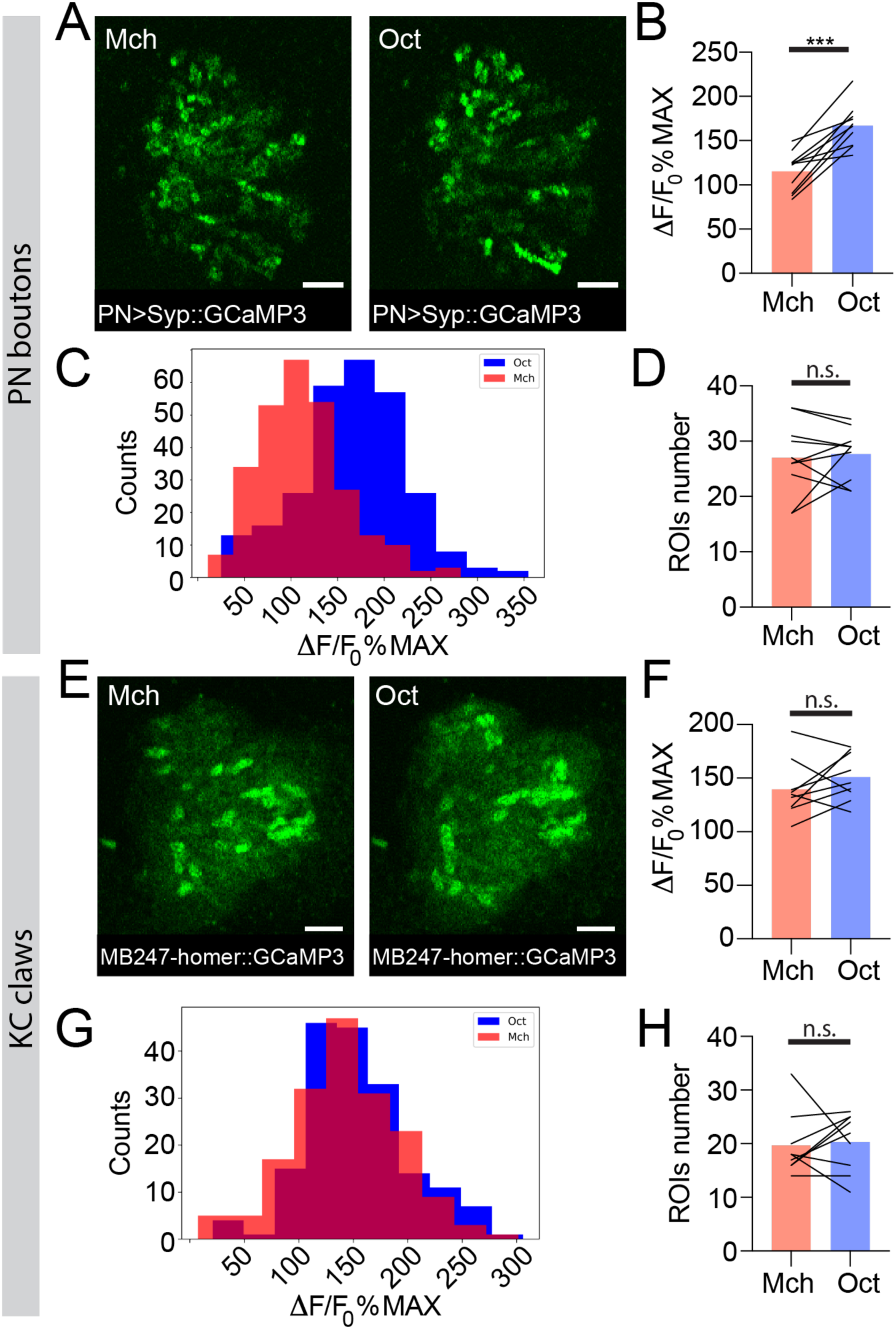
The strength of response to odour stimulation varies in an odour-dependent way in the PN boutons, but is homogenous at the postsynaptic KC claws. **(A)** Example of PN boutons fluorescence increase in response to stimulation with Mch or Oct in *NP225-GAL4>UAS-Syp::GCaMP3* flies. Scale bar = 10μm. **(B)** The average activity peak in PN boutons was higher when flies were exposed to Oct compared to Mch. n=10, p=0.0002, paired t-test. **(C)** Frequency distribution of PN boutons activity peaks in the Mch vs Oct protocol. The Oct population was significantly shifted towards higher *Δ*F/F_0_%MAX values. n=10, p<0.0001, Kolmogorov-Smirnov test. **(D)** The number of ROIs showing odour-evoked activity did not change between the two odour exposures. n=10, p=0.689, paired t-test. **(E)** Example of KC claws fluorescence levels in response to Mch and Oct in *MB247-homer::GCaMP3* flies. Scale bar = 10μm. **(F)** The average activity peak in KC claws was comparable between Mch and Oct exposures. n=9, p=0.1648, paired t-test. **(G)** Frequency distribution of KC claws activity peaks in the Mch vs Oct protocol. The two populations had a similar shape and spread among similar *Δ*F/F_0_%MAX values. n=9, p=0.0982, Kolmogorov-Smirnov test. **(H)** The number of ROIs showing odour-evoked activity did not change between the two odour exposures. n=9, p=0.727, Wilcoxon matched-pairs test. Odours were diluted 1:100, bars indicate means.

To clarify the impact of the odour-tuned activation of APL on the response of KCs to odours, we next imaged the functional response of KC claws to odour stimulation. Flies expressing the postsynaptically-tagged calcium indicator homer::GCaMP3 under the KCs promoter *MB247* (Pech et al. 2015) were prepared, stimulated and imaged as described before. Olfactory stimulation caused the activation of different patterns of MGs in an odour-dependent manner (Pech et al. 2015) (Fig 3E, S3F). The number of MGs responding to each odour was not significantly different between Mch and Oct exposure (Fig 3H, n=9, p=0.727, Wilcoxon matched-pairs test), whereas it was lower when flies were stimulated with ẟ-DL (Fig S3I, n=10, p=0.0059, paired t-test), which elicits the least overall ORN activity (Hallem and Carlson 2006a) and induced a weak and restricted response in PN glomeruli at the AL (Fig S2). Importantly, the average microglomerular postsynaptic response to each odour was not different when comparing the response to Mch vs Oct stimulation (Fig 3F, n=9, p=0.1648, paired t-test) or ẟ-DL vs Oct stimulation (Fig S3G, n=10, p=0.767, Wilcoxon matched-pairs test). Additionally, the frequency distributions of the odour-evoked activity peaks were overlapping (Fig 3G, S3H, n=9-10, p=0.0982 and p=0.9554 for Fig 3G and S3H, respectively, Kolmogorov-Smirnov test). Hence, the differences in activation strength described at the input population of MGs (the PN boutons) seemed to be normalized at the next neuronal layer, in the KC claw-like dendritic endings. Thus, the range of postsynaptic responses in MGs appears to be restrained.

### 2.4 APL silencing leads to more variable odour evoked activity at the MGs postsynapses

Taken together, we showed that APL activation varies with different odours and scales with PN boutons activation levels. Together with the higher similarity among KC claws responses to different odours, this suggests a role of APL as normalizer of olfactory input-elicited response at the KC dendritic claws. If this is correct, blocking APL output would possibly lead to more variable activation of KC claws, mirroring PN bouton activation. We tested this hypothesis by expressing in APL tetanus toxin light chain (TNT), to block vesicle exocytosis and thus silence APL output (Sweeney et al. 1995). To suppress toxin expression during development, we co-expressed the temperature sensitive GAL4 inhibitor *tubP-GAL80^ts^*. Flies were kept at 18 °C until eclosion, and then transferred at 31 °C for 24h to 48h prior to the experiments. KC responses in APL-silenced flies (APL OFF) were imaged with the postsynaptically-tagged *MB247-homer::GCaMP3* construct. Due to the stochastic nature of *APLi-GAL4* mediated expression (A. C. Lin et al. 2014), flies in which the flippase-dependent expression of TNT in APL did not happen were imaged as control (APL ON). Animals were stimulated with Mch or Oct. As expected, control animals neither show a difference in the average peak response to the two stimuli (Fig 4B, n=10, p=0.949, 2-way ANOVA with Tukeýs multiple comparisons) nor in the number of responding units (Fig 4D, n=10, p=0.995, 2-way ANOVA with Tukeýs multiple comparisons). Furthermore, the frequency distributions of the odour-evoked activity peaks overlapped with each other (Fig 4E, n=10, p=0.0533, Kolmogorov-Smirnov test; see also Fig 3G). In calyces where the APL output was blocked by TNT expression though, we measured a significant difference in the response to the two tested odours, with Oct causing a stronger average odour-evoked activity (Fig 4B, n=10, p=0.0003, 2-way ANOVA with Tukeýs multiple comparisons) as well as a slight increase in the number of responding MGs (Fig 4D, n=10, p=0.047, 2-way ANOVA with Tukeýs multiple comparisons). Interestingly, small clusters of responding MGs were often found spatially close to each other in the APL OFF scenarios (e.g. compare spatial distribution of Oct responders in 4C vs 4A) suggesting a possible locality of APL-mediated inhibition. Finally, the frequency distribution plot for the APL OFF flies showed two shifted populations, with Oct responses skewed towards higher values (Fig 4F, n=10, p<0.0001, Kolmogorov-Smirnov test), resembling the distribution displayed by the PN boutons (Fig 3C). In summary, blocking APL output led to a more variable odour representation at the level of KC claws. This variable odour representation bore a higher similarity to the activity of the input population, hence supporting the hypothesis that APL acts as an input strength normalizer on MGs in the MB calyx.

**Fig 4.**
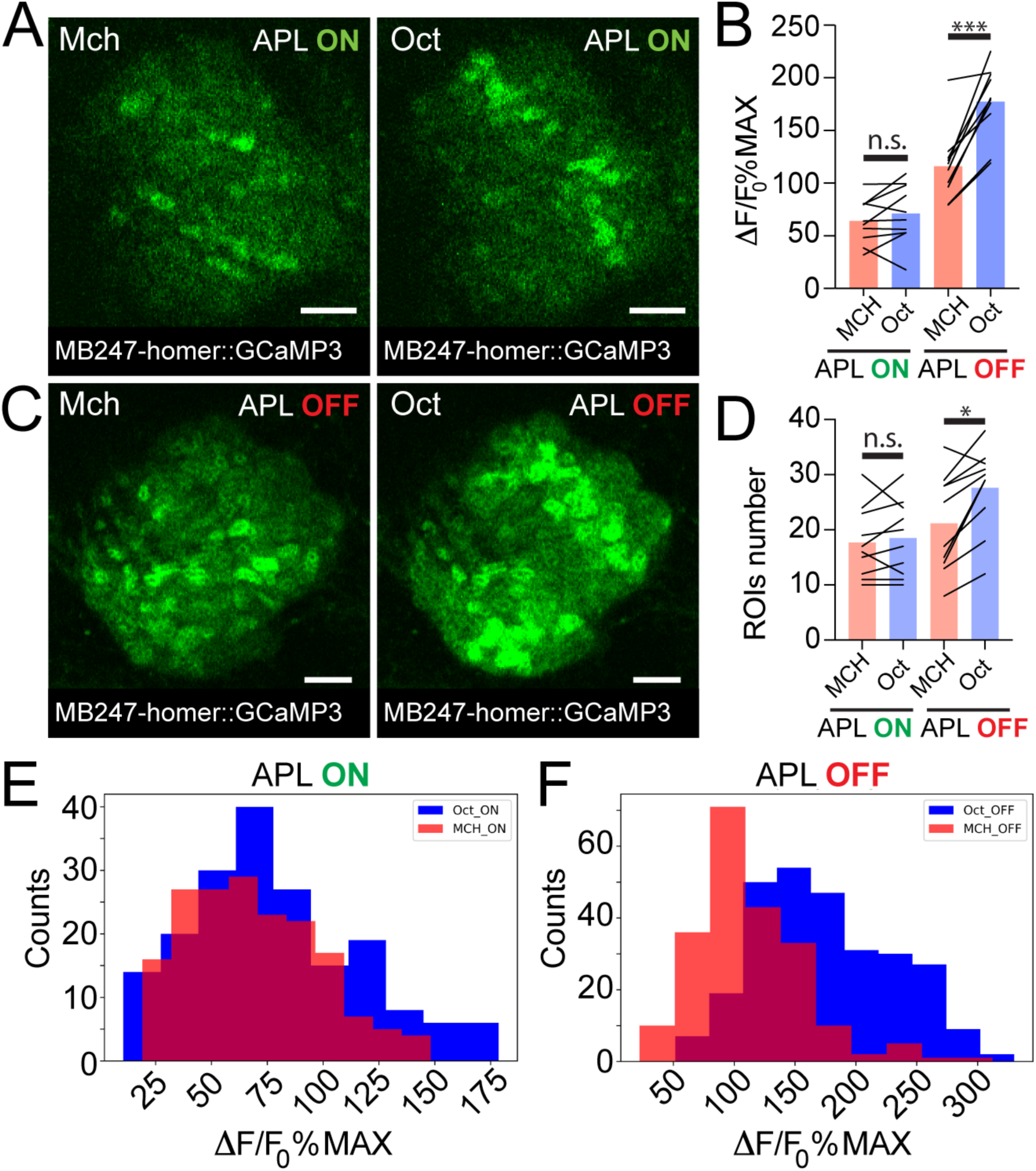
APL silencing leads to more variable odour representations at the MB calyx. Examples of KC claws fluorescence levels in response to Mch and Oct in control animals **(A**, APL ON) or in flies where the output from APL was blocked **(C**, APL OFF). The genotype used is *APLi-GAL4>UAS-TeTx, UAS-mCherry; MB247-homer::GCaMP3*. Scale bar = 10μm. **(B)** The average activity peak in KC claws was similar when APL was active (APL ON, p=0.949), but was highly variable in the absence of APL output (APL OFF, p=0.0003). n=10, 2-way ANOVA with Tukeýs multiple comparisons. **(D)** The number of odour-responding ROIs was comparable in the presence of active APL (APL ON, p=0.995) and it was slightly increased in the absence of APL output (APL OFF, p=0.047). n=10, p=0.047, 2-way ANOVA with Tukeýs multiple comparisons. **(E)** Frequency distribution of MGs activity peaks in the presence of APL activity. The two populations are highly overlapping, as in Fig 3G. n=10, p=0.0533, Kolmogorov-Smirnov test. **(F)** In the absence of APL inhibition, the distribution of MGs responding to Oct shifted towards higher values, resembling presynaptic PN boutons data shown in Fig 3C. n=10, p<0.0001, Kolmogorov-Smirnov test. Odours were diluted 1:100, bars indicate means.

### 2.5 APL inhibition onto MGs of the MB calyx is local

Our data indicate that APL contribution to the MG microcircuit yields a normalized postsynaptic response, independently of the variability of olfactory input strength. To understand how this input-tuned inhibition is achieved, we next asked whether the inhibition provided by APL is global and equally delivered to the entire calyx, or whether it might be more locally restricted to the sites of PN activation. Towards this aim, we selected 3 sets of PNs: DM3, VA1d and DC3, that appear to have distinctive bouton distributions in the calyx based on the Hemibrain dataset (F. Li et al. 2020b). We expect the bouton distribution of these PNs to be reproducible among animals (H.-H. Lin et al. 2007; Jefferis et al. 2007). Based on an available database of odorant representations (Münch and Galizia 2016), these PNs are activated by the following odours: Pentyl acetate (PA) activates the glomerulus DM3, whose PNs terminate in the posterior part of the calyx mainly; Methyl Palmitate (MP) activates the glomerulus VA1d, whose PN terminals populate the anterior part of the calyx; Farnesol (FA) activates glomerulus DC3, with PN terminals that populate the anterior part of the calyx, similarly to VA1d (Fig 5A-B). Flies were exposed to random sequences of the 3 odours, and 3D image stacks were acquired over time to define the localization of the APL response within the entire calycal volume (Fig 5B). Interestingly, we found the ratio of the calcium transients to be highly inconstant throughout the volume of the calyx for the structurally more distant odours PA and MP (see also Material and Methods for details). In particular, the PA/MP fluorescence ratio was higher in the posterior sections of the calyx and reduced in the anterior ones (Fig 5C, black line), reflecting the bouton distribution of the PNs activated by these two odours. By contrast, the fluorescence ratio of FA to MP was more homogeneous across the calyx volume, reflecting the fact that the boutons of the PNs that respond to these odours are located in a similar region of the calyx (Fig 5C, blue line). In other words, not only APL calcium transients were different with different odours, as already shown (Fig 2), but these transients were also differently distributed in the calycal volume depending on the odour, strengthening the link between structural and functional data. Combined with the information about synapse distribution, these data suggest that APL is locally activated in the calyx at the MGs corresponding to activated PNs, and that the tone of APL inhibition in the calyx is a gradient that slowly degrades with increasing distance from the active boutons. Taken together, APL inhibition onto MGs of the MB calyx showed signs of locality. Of notice, a similar spreading mechanism has been observed in APĹs parallel neurites at the MB lobes, where calcium transients failed to propagate over long distances (Amin et al. 2020).

**Fig 5.**
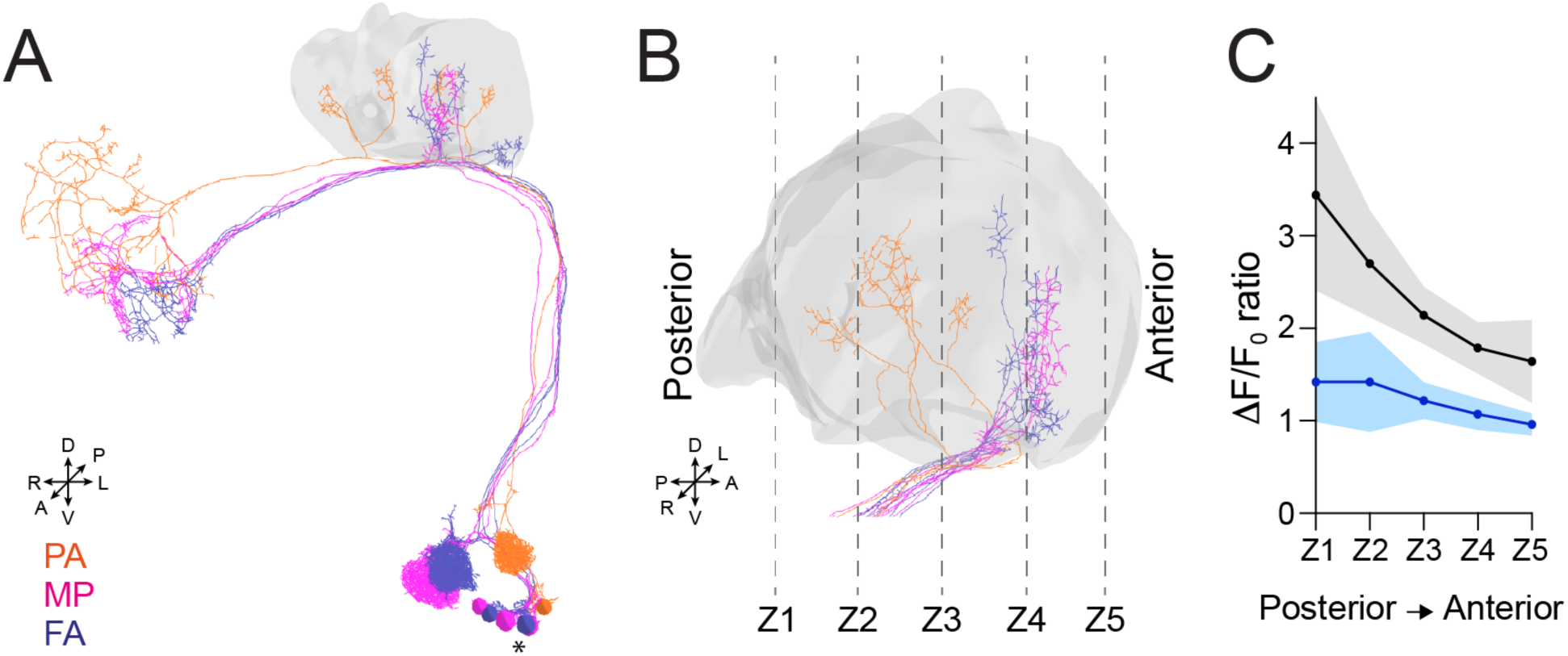
APL inhibition is local within the MB calyx. **(A)** 3D skeletons of the PNs activated by the odours used in this experiment: PA (orange), MP (magenta), FA (blue). Asterisk indicate PN cell bodies. **(B)** Side view of the calyx showing the distribution of the PN terminals for the odours used in this experiment. Note the higher spatial segregation between PA and MP PN terminals compared to FA and MP ones. The ticked lines Z1-Z5 show an example of sectioning applied when acquiring image stacks over time. **(C)** The calcium transient ratio of PA to MP (black line) was highly variable throughout different section of the calyx, and was significantly different to the FA to MP one (blue line). n=7, p=0.007 (**), Kolmogorov-Smirnov test. Coloured areas represent SEM.

## 3 Discussion

While the importance of inhibition in reducing the overlap among stimuli representation has been postulated many decades ago (Marr 1969; Albus 1971; Litwin-Kumar et al. 2017; Cayco-Gajic and Silver 2019) and supported by more recent experimental evidence (Parnas et al. 2013; Olsen, Bhandawat, and Wilson 2010), the complete mechanism by which inhibition supports stimuli discrimination is not fully understood yet. Here, we show that the inhibitory APL neuron, by participating in the structure of MGs of the *Drosophila* MB calyx, provides inhibition scaled to the PNs excitatory inputs to the calyx. As a result, the average strength of postsynaptic responses in KC dendritic claws is homogenous among MGs and different odour representations. We suggest that this normalization of postsynaptic responses operated by APL is at the core of pattern separation in the MB.

Pattern separation is obtained in the MB through the formation of a sparse response in the KC layer (Honegger, Campbell, and Turner 2011; Turner, Bazhenov, and Laurent 2008; Campbell et al. 2013a; Perez-Orive et al. 2002). The decoding of a sparse code, in general, increases the storage capacity of associative networks, thereby supporting learning and classification tasks (Olshausen and Field 2004; Kanerva 1988; Tsodyks and Feigel’man 1988; Perez Vicente and Amit 1989; Huerta et al. 2004; García-Sanchez and Huerta 2003; Jortner, Farivar, and Laurent 2007). In fact, sparse neuronal representations are described in several organisms including mammals, songbirds, and insects (Rolls and Tovee 1995; Vinje and Gallant 2000; Hahnloser, Kozhevnikov, and Fee 2002; Laurent 2002; Quiroga et al. 2005; GoodSmith et al. 2017; Danielson et al. 2017). APL was reported to play a key role in maintaining KCs responses sparse (A. C. Lin et al. 2014; Lei et al. 2013), but the underlying mechanism was far from understood. KCs receive inputs from 6-8 PNs on average (Butcher et al. 2012; Zheng et al. 2020; F. Li et al. 2020b) and, due to KCs high firing threshold (Turner, Bazhenov, and Laurent 2008), require more than half of those inputs to be coactive to spike (Gruntman and Turner 2013). Our data suggest that the APL neuron, by confining KC claws responses within a certain range of activation, ensures that KCs requirement of multiple coactive claws is respected even in the presence of highly variable input strengths. In other words, APL inhibition makes KC input integration dependent on the combinatorial pattern of inputs rather than on the strength of individual inputs. Of notice, odour discrimination is achieved at multiple levels of the *Drosophila* olfactory pathway by different means of inhibitory mechanisms: from the input gain control normalization executed by GABAergic interneurons in the AL (Olsen and Wilson 2008), to the high-pass filter function performed by inhibitory iPNs at the lateral horn (Parnas et al. 2013).

Interestingly, we found a slight increase in the number of MGs responding to an odour in the absence of APL inhibition (Fig 4D). This could have two possible explanations: i) the postsynaptic response of these additional MGs was so weak in the presence of APL that their activation was below the limit of detection; or ii) the increase in number is the consequence of an expanded number of KC axons spiking action potentials that back-propagate into their dendrites. We tend to exclude the second hypothesis as each MG is constituted, on average, by 13 claw-like dendritic endings from different KCs (Davi D. Bock, personal communication), and each KC connects with its other claws to random PNs (Caron et al. 2013). It seems unlikely that all or a large fraction of randomly assembled claws are all invested by back-propagating action potentials and therefore visible in our recordings.

Our structural and functional data point towards the involvement of APL in a feedforward loop from PN boutons to KC claws, as well as a closed feedback loop with PN boutons. An advantage of using recurrent circuits to provide inhibition is that such a system can deal with a wide range of input strength, as inhibition and excitation strengths are proportional. Indeed, EM analysis revealed both pre- and postsynaptic connection between APL and PN boutons, linearly proportional to each other (Fig 1D), and the differences in the APL calcium influx in response to odours correlated to the variability measured in PNs (Fig 2 and 3). So far, APL has been mainly described as a feedback neuron for KCs (A. C. Lin et al. 2014; Lei et al. 2013; Amin et al. 2020). However, feedforward inhibitory neurons from the input population onto the next layer have been described in other neuronal networks performing pattern separation (Cayco-Gajic and Silver 2019). For example, granule cells receive both feedforward and feedback inhibition from Golgi cells at the cerebellar cortex (Vos, Volny-Luraghi, and de Schutter 1999; Duguid et al. 2015), which are driven by excitatory inputs from the mossy fibers (Kanichay and Silver 2008) and granule cells’ axons, respectively (Cesana et al. 2013). Moreover, it has recently been demonstrated that Golgi cells recruitment scales with the mossy fibers input density (Tabuchi et al. 2019), similarly to what we observed in our functional imaging experiments. Additionally, adaptive regulation of KCs sparseness by feedforward inhibition has already been theorized in realistic computational models of insect’s mushroom bodies (Assisi et al. 2007; Luo, Axel, and Abbott 2010). Regarding KCs-to-APL connections, we found a positive linearity among pre- and post-synapses between these two cells (Fig 1E), confirming the presence of a local feedback loop within KC dendrites and the APL at the calyx (Amin et al. 2020). Furthermore, we reported that *α*/*β* KCs receive more inhibitory synapses along their dendritic trees compared to *γ* and *α*’/*β*’, where the majority of synapses received from the APL is localised on KC claws instead (Fig S1C). As the ability of inhibitory synapses to shunt current from excitatory synapses depends on the spatial arrangement of the two inputs (Spruston, Stuart, and Häusser 2016), we speculate that the difference in APL synapses localisation could contribute to some of the electrophysiological differences recorded among distinct KCs type. For example, *α*/*β*_c_ KCs were found to have a higher input resistances and longer membrane time constants compared to *α*’/*β*’ KCs (Groschner et al. 2018), resulting in a sigmoidal current-spike frequency function rather than a linear one (Groschner et al. 2018). Additionally, a difference in synapses distribution can also indicate that two inhibitory mechanisms coexist at the MB calyx, similarly to what has been shown in the cerebellum where Golgi cells are responsible for both tonic inhibition, controlling granule cells spike number (Brickley, Cull-Candy, and Farrant 1996), gain control (Mitchell and Silver 2003), and phasic inhibition, limiting the duration of granule cells responses (D’Angelo and de Zeeuw 2009).

Finally, volumetric calcium imaging showed that the APL inhibition is local within the MB calyx. In particular, we found a difference in the APL calcium transients when flies were stimulated with odours that activate PN subsets with segregated bouton distribution in the calyx (Fig 5). These data suggest that APL inhibition onto MGs can be imagined as a gradient that peaks at the MGs active during a given stimulus and attenuates with distance. Non-spiking interneurons in insects are typically large and characterized by complex neurite branching, an ideal structure to support local microcircuits (Roberts, Bush, and Society for Experimental Biology (Great Britain). Neurobiology Group. 1981). As a matter of fact, similar examples of localized APL response as described here have been reported in the *Drosophila* MB (Amin et al. 2020; Wang et al. 2019) as well as in the APĹs homolog GGN in the locust (Ray, Aldworth, and Stopfer 2020; Leitch and Laurent 1996). An advantage of having local microcircuits is that it allows a single neuron to mimic the activity of several inhibitory interneurons, as described in amacrine cells of both mammals (Grimes et al. 2010) and *Drosophila* (Meier and Borst 2019). Additionally, a parallel local-global inhibition is suggested to expand the dynamic range of inputs able to activate KCs (Ray, Aldworth, and Stopfer 2020).

An important open question is whether the APL inhibition onto MGs of the MB calyx is more of a presynaptic phenomenon, therefore acting on PN boutons output, or a postsynaptic one on KCs claws. Our functional data reveal a clear impact of APL on the postsynaptic response in MGs (Fig 3H, S3F), while the PN boutons display a broad range of activity levels (Fig 3A-D, Fig S2, Fig S3A-D). Accordingly, silencing of the GABA_A_ receptor *Rdl* on KCs increased calcium responses in the MB, including the calyx (Liu, Krause, and Davis 2007), and reduced sparseness of odour representations (Lei et al. 2013). However, due to the presence of presynapses from APL to PN boutons (Fig 1B, 1D), a presynaptic component of APL inhibition is certainly possible.

One possible caveat to our hypothesis is given by the fact that reducing GABA synthesis in APL by RNAi has been found to improve olfactory learning (Liu and Davis 2009). However, this could be explained by a low efficiency of RNAi in this case. Indeed, incomplete silencing might increase KCś output without affecting sparseness. As a matter of fact, blockage of APL output via *shibire^ts^* led to impaired olfactory discrimination (A. C. Lin et al. 2014).

Taken together, our study provides novel insights on how feed-forward inhibition via APL shapes the postsynaptic response to olfactory inputs in the MB calyx and contributes to maintaining odour evoked KC activity sparse. In the future, it will be interesting to investigate the impact of APL on memory consolidation, which has been associated with structural plasticity in the calyx (Baltruschat et al. 2021) and with changes in the KC response (Delestro et al. 2020).

## 4 Materials and Methods

### 4.1 Connectomics

Connectomics data were obtained from the Hemibrain EM dataset (Scheffer et al. 2020) via the Neuprint analysis Tool (Clements et al. 2020). In particular, the Neuprint-python package (https://github.com/connectome-neuprint/neuprint-python) was used to filter for annotated synapses made and received by the APL only within the CA(R) ROI. The command *fetch_adjacencie*s was used to extract data regarding the connectivity among cell types. To visualize neuron skeletons or filled renders in 3D, the commands *fetch_skeletons* or *fetch_mesh_neuron* were used instead. To visualize APL synapses onto PNs or KCs, the coordinates of those synapses were obtained via *fetch_synapse_connections* and plotted together with the 3D neuronal meshes. The localization of the synapses (e.g., on PN bouton or not) was addressed manually by two separate users in a blind manner, and the average counts were calculated. Detailed tables containing the list of all PNs and KCs interconnected with the APL within the MB calyx, as well as the weight of those synapses, and 3D images of APL synapses mapped onto PNs and KCs meshes can be found at: Mendeley data link will be available upon publication.

### 4.2 Fly strains

The following lines were used for experiments: *GH146-Gal4* (Reinhard F. Stocker et al. 1997), *NP225-Gal4* (Hayashi et al. 2002), *NP2631-Gal4* (Hayashi et al. 2002), *GH146-Flp* (Hong et al. 2009), *tubP-GAL80^ts^* (Sean E. McGuire et al. 2003), *tubP-FRT-GAL80-FRT* (S. Gao et al. 2008; Gordon and Scott 2009), *UAS-GCaMP6m* (Chen et al. 2013a), *MB247-homer::GCaMP3* (Pech et al. 2015), *UAS-Syp::GCaMP3* (Pech et al. 2015), *UAS-TeTx* (Sweeney et al. 1995)*, UAS-mCherry∷CAAX* (Kakihara et al. 2008). Flies were raised in a 12h/12h light-dark cycle on a standard cornmeal-based diet at 25 °C, 60% relative humidity unless they expressed the temperature-sensitive gene product Gal80^ts^. Flies carrying *tubP-GAL80^ts^* were raised at 18°C and placed at 31°C for 24 h-48 h <24 h after eclosion. 1-7 days old flies were used for experiments. All experiments were performed on mixed populations of males and females.

### 4.3 Two-photon *in vivo* calcium imaging

For *in vivo* imaging in the MB calyx, adult flies were briefly anaesthetized on ice, positioned in a polycarbonate imaging chamber (Louis et al. 2017), and immobilized using Myristic Acid (Sigma-Aldrich). To allow optical access to the Calyx, a small window was opened through the head capsule under Ringer’s solution (5 mM HEPES, pH 7.4, 130 mM NaCl, 5 mM KCl, 2 mM CaCl_2_, 2 mM MgCl_2_; pH adjusted to 7.2). To minimize movement, fly heads were stabilized with 1,5% low melting agarose (Thermo Scientific) in Ringer’s solution, immediately before dissection. Flies were imaged with a two-photon laser-scanning microscope (LaVision BioTec, TriM Scope II) equipped with an ultra-fast z-motor (PIFOC® Objective Scanner Systems 100µm range) and a Nikon 25X CFI APO LWD Objective, 1.1 NA water–immersion objective. GCaMP molecules were excited at 920 nm using a Ti:sapphire laser (Coherent Chameleon). Odours were delivered to the *in vivo* preparation via a 220A Olfactometer (Aurora Scientific) in a randomized fashion. Odours were loaded into the respective odour vials with a dilution 10X higher than the desired one, and further diluted 1:10 with clean air during odour stimulation. A constant flow of clean air was provided by a Stimulus Controller CS 55 (Ockenfels SYNTECH GbmH), equipped with two activated carbon inlet filters to avoid air contamination. Animals were stimulated with 2 odour puffs of 5 s each, separated by 20 s clean air intervals. Both clean air and odour flows were kept around 0.5L/min for the entire experimental procedure. For imaging, a region large enough to include an entire z-section of the mushroom body calyx was chosen. The scanning frequency was set around 9 Hz. Single plane videos were acquired unless stated otherwise.

For volumetric calcium imaging (Fig 5), flies were mounted and stimulated as described above and 3D stacks of 5 images each were acquired over time for the entire stimulation time. To compensate for the reduced speed caused by the stack acquisition, the frame rate was adjusted to around 16 Hz.

For *in vivo* AL imaging, female adult flies were briefly anaesthetized on ice, positioned in a custom-built fly chamber (Hancock, Bilz, and Fiala 2019), and immobilized using UV-hardening dental glue (Kentoflow, Kent Dental). A small dissection was performed in Ringer’s solution (5 mM HEPES, pH 7.4, 130 mM NaCl, 5 mM KCl, 2 mM CaCl_2_, 2 mM MgCl_2_; pH adjusted to 7.2). Flies were imaged with a two-photon laser-scanning microscope LSM 7MP (Zeiss) equipped with a 20X/1.0 DIC M27 75mm Plan-Apochromat objective (Zeiss). GCaMP molecules were excited at 920 nm using a Ti:sapphire laser (Coherent Chameleon). Odours were delivered to the *in vivo* preparation via a custom-built olfactometer. Odours were diluted in mineral oil (Sigma-Aldrich) at the required dilution. A constant flow of clean air was provided by a membrane pump “optimal” (SCHEGO). Animals were stimulated with 2 odour puffs of 5 s each, separated by 20 s clean air intervals. Both clean air and odour flows were kept around 1mL/s for the entire experimental procedure. For imaging, a region large enough to include an entire z-section of the mushroom body calyx was chosen. The scanning frequency was set around 5 Hz. Single plane videos were acquired unless stated otherwise.

As the PN Gal4 driver *GH146* used in AL imaging experiments drives expression also in APL (Liu and Davis 2009), the *NP225-Gal4* driver line (Hayashi et al. 2002) was chosen for PN boutons imaging at the calyx, as it targets a similar amount of PNs as *GH146* without including the APL.

### 4.4 Data Analysis

Two-photon images were analysed using Fiji (Schindelin et al. 2012). Briefly, raw videos were motion corrected via the “Template Matching and Slice Alignment” ImageJ plugin (Tseng et al. 2012). Afterwards, ROIs of the single MGs responding to a given odour were created via the “Cell Tracking by Calcium” ImageJ Macro (designed and written by DZNE IDAF). The generated mask was then applied to the previously registered video in order to extract the average intensity value over time per each of the detected ROIs. Finally, the ΔF/F_0_% and the ΔF/F_0_%MAX of each ROI was calculated by using the average intensity of the first 30 frames as F_0_. A detailed manual related to the ImageJ Macro, as well as Python notebooks computing the ΔF/F_0_% and ΔF/F_0_%MAX given a dataframe of intensity values over time, is available at: Mendeley data link will be available upon publication.

To measure calcium influx among the APL neurites branching in the mushroom body calyx, a manual ROI was drawn around the entire calycal region expressing the GCaMP and the average intensity value over time was extracted. F/F_0_% and ΔF/F_0_%MAX were calculated as described above.

For the odours response ratio calculated in the APL volumetric calcium imaging experiment (Fig 5), the ΔF/F_0_%MAX per each odour was calculated as described above. The ΔF/F_0_%MAX values per each of the 5 frames contained in an image-stack were obtained and averaged among all animals tested. Next, a ΔF/F_0_%MAX ratio between the odour pairs being compared was calculated and analysed per each of the sections included in an image stack.

### 4.5 Confocal imaging

To address the presence or not of the *APLi.GAL4* driven *UAS-TNT* and *UAS-mCherry* products, whole flies were fixed on Formaldehyde (FA) 4% in PBS with 0.1% Triton X-100 (PBT 0.1%) immediately after *in vivo* imaging. Once all animals sustained the *in vivo* imaging protocol, brains were dissected using a pair of forceps in a small petri dish covered with a layer of silicon, fixed for further 20 min on FA 4% in PBT 0.1%, washed 3 times for 5 min ca in PBT 0.1% and mounted with Vectashield® Plus Antifade Mounting Medium (Vectorlabs) on 76 x 26 mm Microscope slides (Thermo scientific) with 1# coverslips (Carl Roth). Brains were oriented with the dorsal part facing upwards. Imaging was performed on a LSM 780 confocal microscope (Zeiss) equipped with a Plan-Apochromat 63x/1.4 Oil DIC M27 objective (Zeiss). 512×512 pixels images were acquired, covering a region of the brain big enough to include the APL soma and branches around the MB calyx and MB lobes. Brains and their related *in vivo* imaging data were assigned to the classes “APL OFF” or “APL ON” based on the presence or not of the TNT-co-expressed mCherry fluorescence, respectively.

### 4.6 Statistics

Statistics were carried out in Prism 8 (GraphPad). Parametric (t-test, ANOVA) or non-parametric tests (Wilcoxon, Mann-Whitney, Kruskal-Wallis, Kolmogorov-Smirnov) were used depending on whether data passed the D’Agostino-Pearson normality test. Statistical power analysis was conducted in G*Power (Faul et al. 2007).

## Acknowledgements

We thank LMF and IDAF sections at DZNE, Seth Tomchick, Sanjeev Kaushalya and Kevin Kuepper for support in technical development. Rita Kerpen, Phuong Tran and Olga Sharma for technical assistance. We thank the Bloomington Stock Center, Andrew Lin and Oren Schuldiner for fly lines. We thank Andrew Lin, Martin Nawrot, Peter Kloppenburg for discussions. We thank Moshe Parnas, David Owald, Barbara Schaffran and the members of the Tavosanis lab for critical reading of the manuscript. This work was supported by the DFG FOR 2705 to G.T.

## 5 Financial interests or conflicts of interest

The authors declare no competing interests.

---

**Fig S1 (Related to Fig 1).**
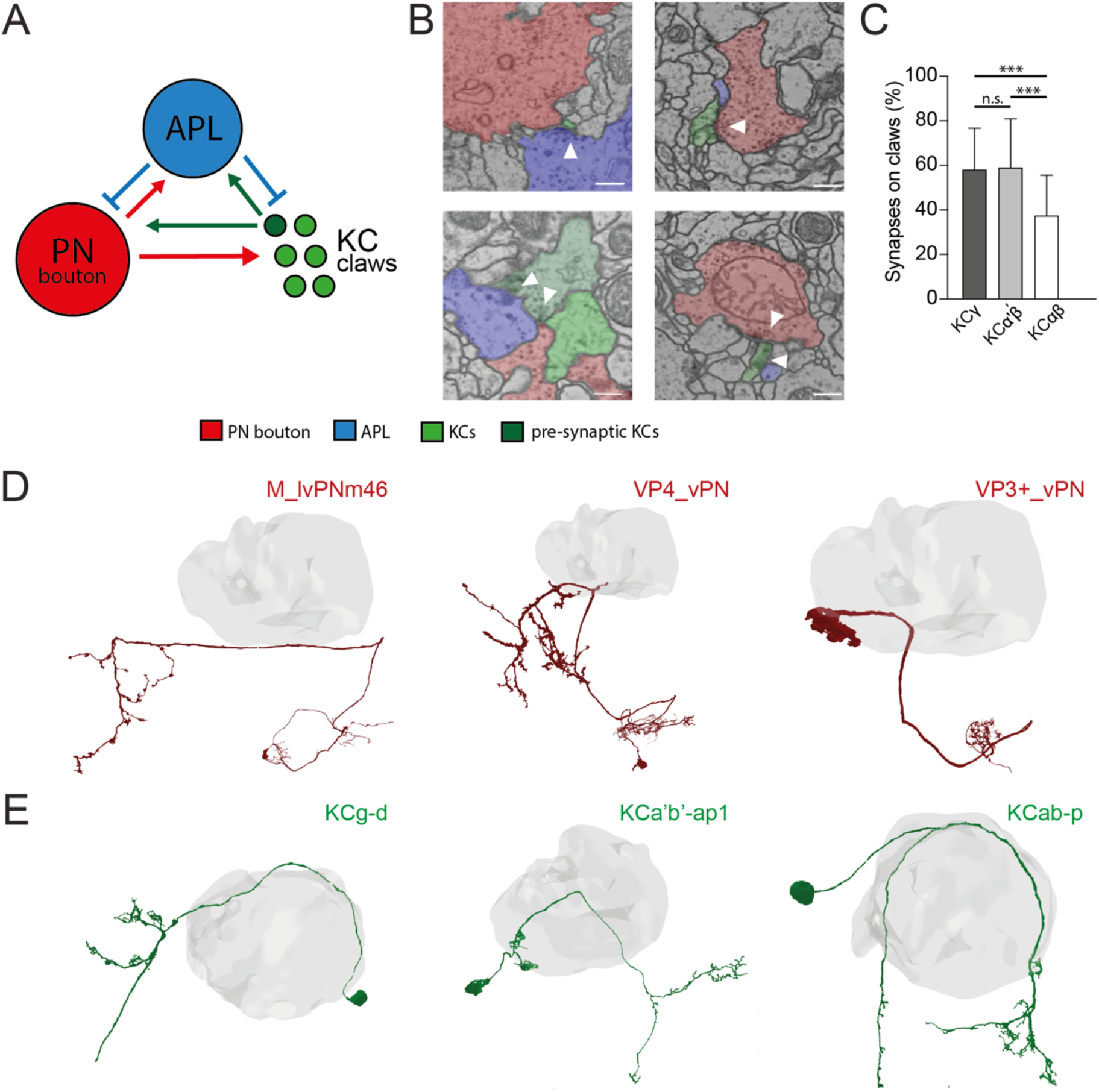
APL in the microglomerular circuit. **(A)** Schematized view of the connectivity patterns found in the DA1-PN MG reconstructed in Baltruschat et al. (2021). Green circles represent KC claws. Dark green circles represent presynaptic KCs. The red circle represents the reconstructed bouton of a DA1-PN. The blue circle represents APL. See Baltruschat et al., (2021) for further explanation, including the full connectome of this DA1-PN MG. **(B)** Examples of connectivity patterns involving APL in the DA1-PN microglomerulus reconstructed in Baltruschat et al. (2021). Top left: EM image of APL (blue) presynaptic to the PN bouton (red) and the claw of a KC (green). Top right: PN bouton presynaptic to APL and three KC claws. Bottom-left: KC (dark green) presynaptic to PN bouton, APL and another claw. Bottom-right: APL and PN bouton presynaptic to the same KC claw. Scale bar = 250nm. **(C)** Spatial distribution of APL synapses among different KC types. The fraction of APL to KC*αβ* synapses localised on KC claws was lower than in other KC types. n=210 (70 per KC type, randomly selected), p<0.0001, Unpaired ANOVA with multiple comparisons. Whiskers indicate SD. **(D)** Examples of PNs not interacting with APL at the MB calyx (grey volume). **(E)** Examples of KCs not interacting with APL at the MB calyx (grey volume). 3D neuronal meshes were created via the Neuprint-python package (see Materials & Methods).

**Fig S2 (Related to Fig 3).**
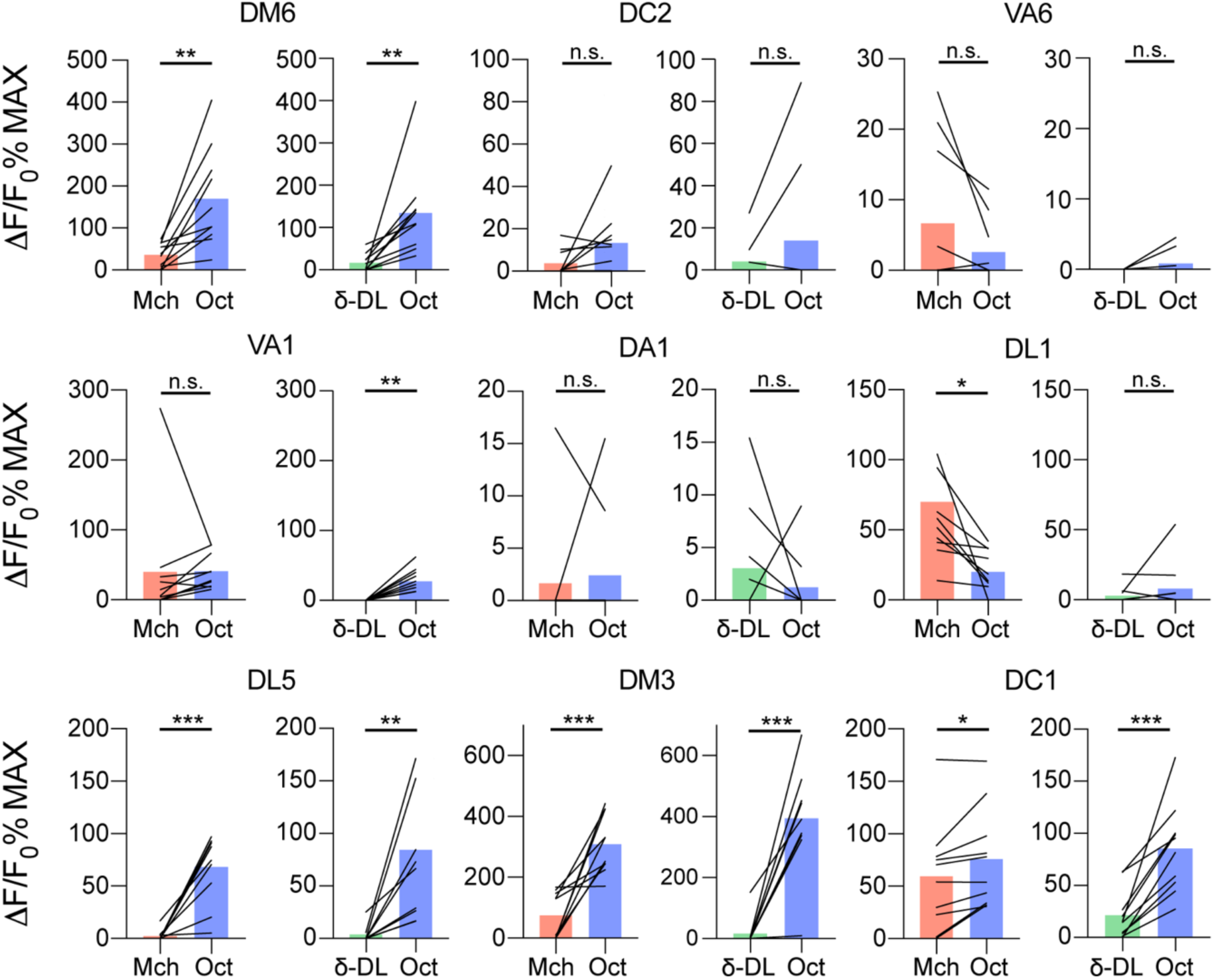
Single PN glomeruli responses at the AL. *GH146-GAL4>UAS-GCaMP6m* flies were stimulated with either Mch vs Oct or ẟ- DL vs Oct. The activity peak per each glomerulus was extracted and compared. n=10, paired t-test, p value > 0.05 (n.s), *≤* 0,05 (*), *≤* 0,01 (**), *≤* 0,001 (***). Odours were diluted 1:100, bars indicate means.

**Fig S3 (Related to Fig 3).**
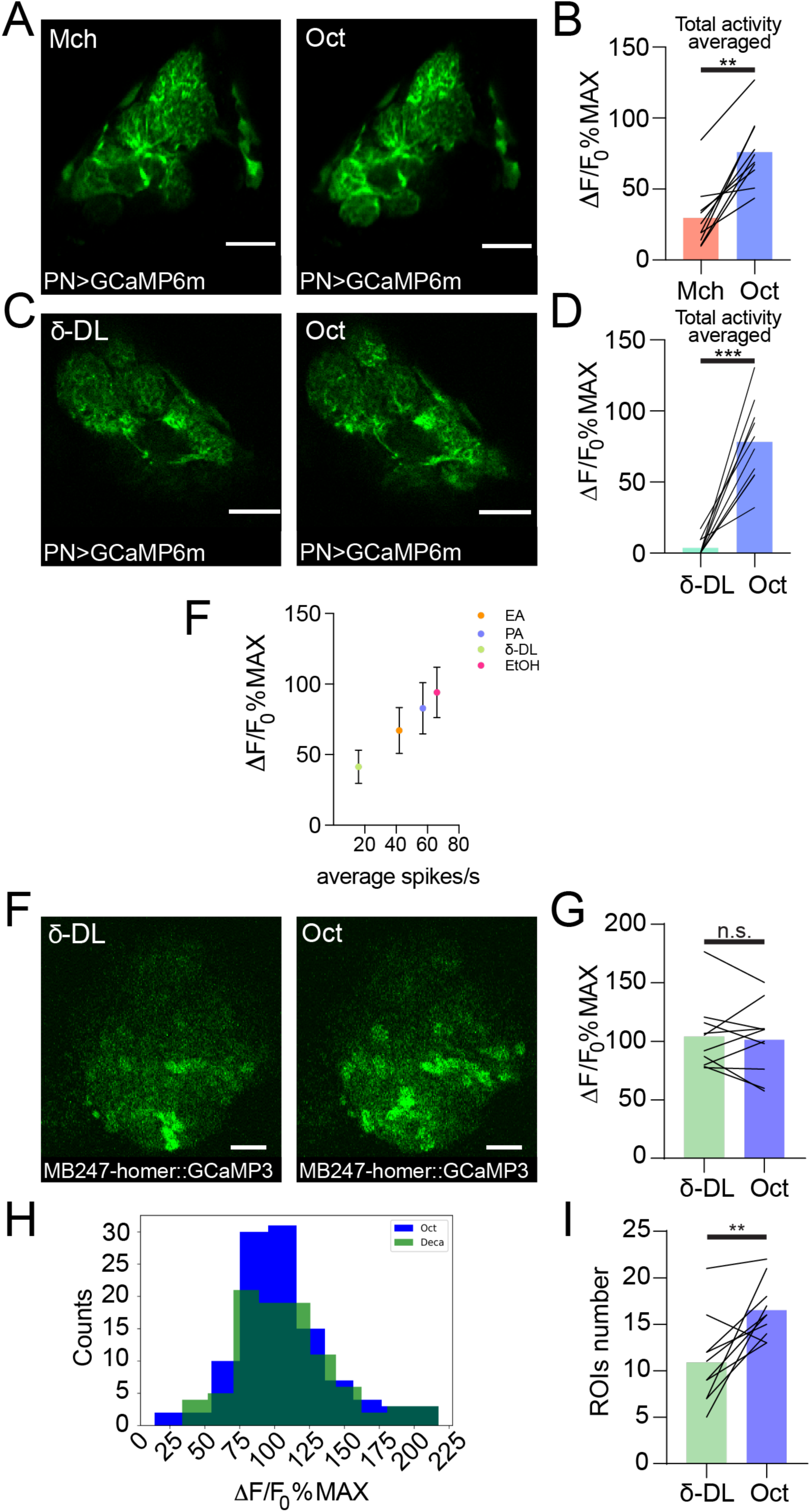
Additional data on PNs and KCs odour evoked activity. Examples of PN dendritic calcium transients in response to either Mch and Oct **(A)** or ẟ-DL and Oct **(C)** at the antennal lobe in *GH146-GAL4>UAS-GCaMP6m* flies. Scale bar = 20μm. **(B)** The average activity peak among responding glomeruli was higher when flies were exposed to Oct compared to Mch. n=10, p=0.002, Wilcoxon matched-pairs test. **(D)** The average activity peak among responding glomeruli was higher when flies were exposed to Oct compared to ẟ-DL. n=10, p<0.0001, paired t-test. **(E)** The APL calcium transient levels in response to 4 chosen odours (ethyl acetate (EA) 1:1000 dilution, pentyl acetate (PA) 1:1000, ẟ-DL and EtOH 1:100) in *APLi-GAL4>UAS-GCaMP6m* flies linearly correlated with the average PN spiking rate for the respective odour stimulations. The PN spiking rate for 24 AL glomeruli were obtained by feeding ORNs spike values from the Hallem and Carlson 2006 dataset into the ORN-to-PN spike transformation equation described in Olsen et al. 2010 (see also (Parnas et al. 2013)). Pearson r=0.99 **(F)** Example of KC claws fluorescence levels in response to ẟ-DL and Oct in *MB247-homer::GCaMP3* flies. Scale bar = 10μm. **(F)** The average activity peak in KC claws was comparable between ẟ-DL and Oct exposures. n=10, p=0.767, Wilcoxon matched-pairs test. **(G)** Frequency distribution of KC claws activity peaks in the ẟ-DL vs Oct protocol. The two populations had a similar shape and spread among similar *Δ*F/F_0_%MAX values. n=10, p=0.9554, Kolmogorov-Smirnov test. **(H)** The number of ROIs responding to Oct was higher compared to the ẟ-DL ones. n=10, p=0.0059, paired t-test. Odours were diluted 1:100 unless stated otherwise, bars indicate means.

